# Preclinical Pharmacokinetics and *In Vitro* Properties of GS-441524, A Potential Oral Drug Candidate for COVID-19 Treatment

**DOI:** 10.1101/2022.02.07.478848

**Authors:** Amy Q. Wang, Natalie R. Hagen, Elias C. Padilha, Mengbi Yang, Pranav Shah, Catherine Z. Chen, Wenwei Huang, Pramod Terse, Philip Sanderson, Wei Zheng, Xin Xu

## Abstract

Preclinical pharmacokinetics (PK) and *in vitro* ADME properties of GS-441524, a potential oral agent for the treatment of Covid-19, were studied. GS-441524 was stable *in vitro* in liver microsomes, cytosols, and hepatocytes of mice, rats, monkeys, dogs, and humans. The plasma free fractions of GS-441524 were 62-78% across all studied species. The *in vitro* transporter study results showed that GS-441524 was a substrate of MDR1, BCRP, CNT3, ENT1, and ENT2; but not a substrate of CNT1, CNT2, and ENT4. GS-441524 had a low to moderate plasma clearance (CLp), ranging from 4.1 mL/min/kg in dogs to 26 mL/min/kg in mice; the steady state volume distribution (Vd_ss_) ranged from 0.9 L/kg in dogs to 2.2 L/kg in mice after IV administration. Urinary excretion appeared to be the major elimination process for GS-441524. Following oral administration, the oral bioavailability was 8.3% in monkeys, 33% in rats, 39% in mice, and 85% in dogs. The PK and ADME properties of GS-441524 support its further development as an oral drug candidate.

## INTRODUCTION

Coronavirus Disease-2019 (COVID-19) caused by severe acute respiratory syndrome coronavirus 2 (SARS-CoV-2) became a global pandemic shortly after its emergence in late 2019. The research community and health care systems have quickly responded to the emergency of the pandemic with vaccine and therapeutics development. Among the successful repurposing attempts, remdesivir (GS-5734), originally developed for Ebola virus [1], has been approved for the treatment of hospitalized COVID-19 patients by the US FDA and other regulatory agencies, after it was shown to reduce the time to recovery by 33% [2]. More recently, remdesivir, given as an early three-day course in mild to moderate COVID-19 patients at risk of progression to severe disease, was shown to reduce progression to hospitalization or death by 87% [3]. However, remdesivir can only be given by intravenous administration (IV) [4] which significantly limits its use [5].

Currently, three COVID-19 vaccines have been approved by US FDA and several other vaccines are under late-stage clinical development. Vaccine development for COVID-19 has progressed at unprecedented speed and resulted in effective vaccines within one year. However, on-going viral gene mutations have been giving rise to new variants of SARS-CoV-2, creating potential antigenic drift and vaccine escape. Therefore, safe, effective, widely available, and orally administered antiviral drugs are still needed for COVID-19 treatment.

Two oral drugs have recently been authorized by the FDA for emergency use for early treatment of mild to moderate COVID-19 patients at high risk of progression to severe disease. Firstly, Paxlovid [6], a combination of nirmatrelvir, an inhibitor of SARS-CoV-2 3C-like protease (3CLpro), and the cytochrome P450 3A4 inhibitor ritonavir, was authorized after it was shown to reduce progression to hospitalization or death by 88% compared to placebo. Secondly, Molnupiravir (MK-4482, EIDD-2801), an oral prodrug of nucleoside N-hydroxycytidine (NHC, EIDD-1931) [7], was authorized after it reduced the progression to hospitalization of death by 30% compared to placebo. Both drugs have limitations: drug-drug interaction potential with Paxlovid, and poor efficacy and mutagenic potential with Molnupiravir. Consequently, there remains a need for effective and safe oral drugs.

Remdesivir is a phosphoramidate prodrug of nucleoside analog GS-441524. GS-441524 is itself a prodrug of the common active compound GS-441524 triphosphate (GS-443902). GS-443902 functions as a pseudosubstrate of SARS-CoV-2 RNA dependent RNA polymerase (RdRP), an enzyme essential for viral RNA replication in host cells. GS-443902 competes with adenosine triphosphate (ATP) as an RdRP substrate, and once incorporated into viral RNA, inhibits RdRP activity [8]. EIDD-1931 triphosphate is also an RdRP substrate, but its mechanism is different, as it induces mutations in the virus RNA leading to error catastrophe. Activation of GS-441524 requires an initial phosphorylation to GS-441524 monophosphate and this step is perceived to be rate limiting in its metabolism to GS-443902. Remdesivir was developed to bypass the initial phosphorylation by using a cell permeable phosphoramidate prodrug which is enzymatically hydrolyzed directly to the monophosphate. As a consequence of the prodrug design, remdesivir has a lengthy synthesis [8]. Remdesivir was studied in a clinical trial to treat the Ebola virus infection in 2015, but its development was terminated owing to lower efficacy compared to antibody-based therapy [9]. GS-441524 is reported to be an effective treatment of feline infectious peritonitis (FIP) caused by feline coronavirus [10]. The cellular uptake and phosphorylation of GS-441524 [11], PK of ester prodrugs of GS-441524 in mice [12], and the *in vivo* efficacy in a SARS-CoV-2 mouse model [13] of GS-441524 have been reported recently. In addition, PK and efficacy of a triester prodrug of GS-441524 in mouse and ferret models of SARS-CoV-2 infection have been reported [14, 15]. The pharmacodynamic study of remdesivir in nonhuman primates exhibited evidence of its conversion to GS-441524 in plasma [16]. Recent clinical reports indicated that GS-441524 was the major circulating metabolite after remdesivir IV administration [17, 18]. Therefore, GS-441524 may have a potential to be further developed as an oral drug for COVID-19 treatment.

In this study, we assessed the oral bioavailability of GS-441524 in animal species and systematically evaluated *in vitro* and *in vivo* ADME (absorption, distribution, metabolism, and elimination) properties of GS-441524, including interactions with nucleoside transporters. PK studies of GS-441524 were performed after IV and PO administrations in C57BL/6 mice, Sprague-Dawley (SD) rats, Cynomolgus monkeys, and Beagle dogs to understand the bioavailability and disposition parameters of GS-441524. All PK data were presented to the public immediately after they were available (https://opendata.ncats.nih.gov/covid19/GS-441524, October 2020). Allometric scaling was utilized to estimate human plasma clearance (CLp) and volume of distribution at steady state (Vd_ss_) from the preclinical PK results. Additionally, the inhibitory activity of GS-441524 was confirmed in Vero E6 cells (derived from the kidney of an African green monkey) and a 3D human airway epithelial (HAE) tissue model of SARS-CoV-2 infection.

## RESULTS

### Inhibition of SARS-CoV-2 infections in Vero E6 cells and a 3D HAE tissue model by GS-441524

To confirm the inhibitory activity of GS-441524 against SARS-CoV-2 infection, we measured the cytopathic effect (CPE) in Vero E6 assay and viral titer in a 3D HAE model of SARS-CoV-2 infection using the methods previously described [19]. GS-441524 inhibited the CPE of SARS-CoV-2 infection in a concentration-dependent manner in Vero E6 cells. The EC_50_ was 1.86 μM, four-fold more potent than that of remdesivir (EC_50_ = 7.43 μM). Neither compound showed any cytotoxicity at 30 μM, the highest concentration tested (Fig. 1A). GS-441524 also reduced viral titer in the EpiAirway 3D HAE tissue model, which contains three pseudostratified layers, including an apical ciliated surface, the underlying mucociliary epithelium, and a basal membrane, and is cultured at air-liquid interface (ALI). The median tissue culture infectious dose (TCID50) was measured after 24 h (Fig. 1B) and 96 h (Fig. 1C) of the initial SARS-CoV-2 infections; GS-441524 significantly inhibited the viral infection, with TCID50 similar to that of remdesivir. Both compounds did not exhibit any cytotoxicity in this model. The results confirmed that the *in vitro* inhibitory activity of GS-441524 against SARS-CoV-2 is similar to that of remdesivir in the 3D HAE tissue model but slightly more potent in Vero E6 cells.

**Figure 1.**
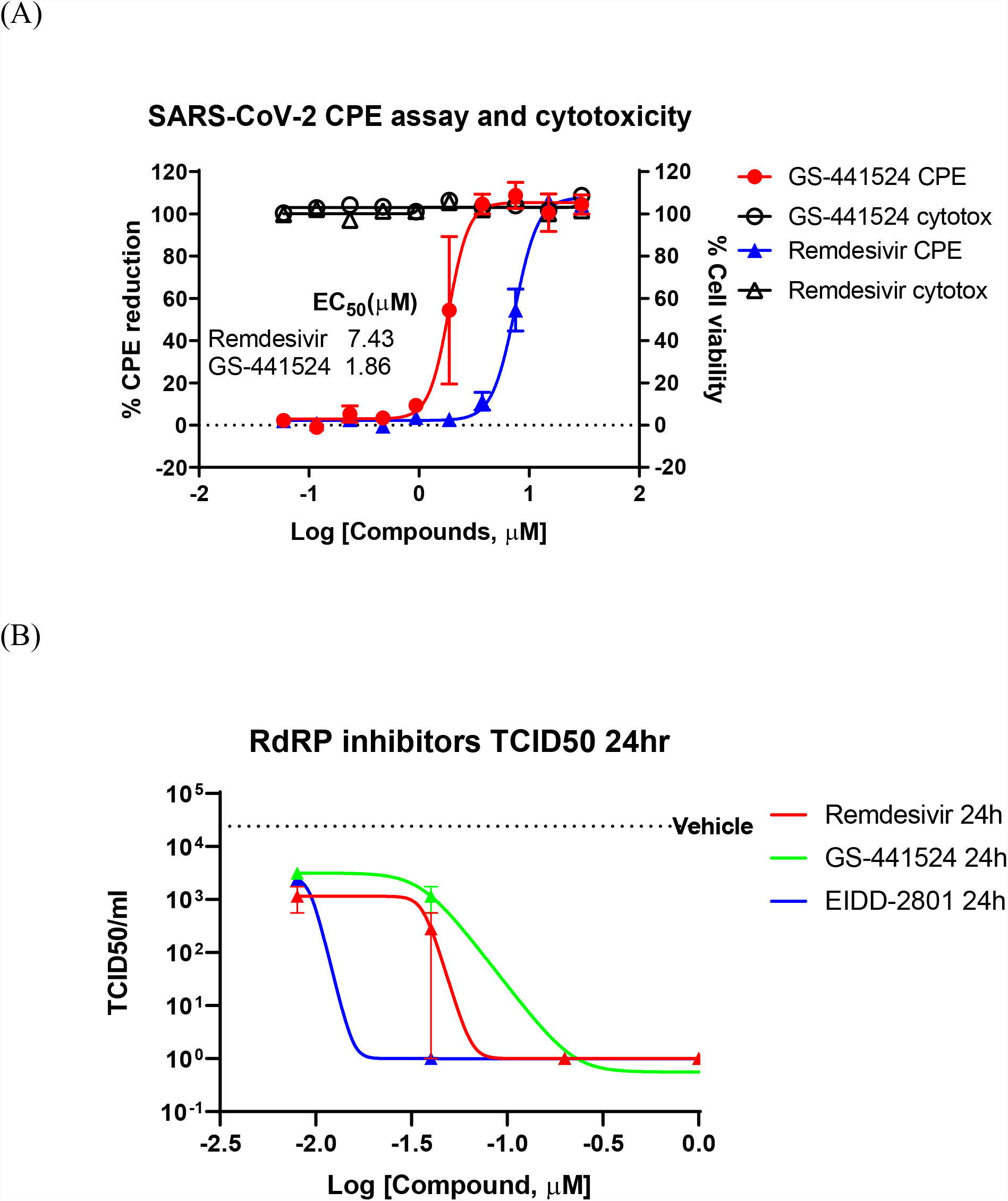
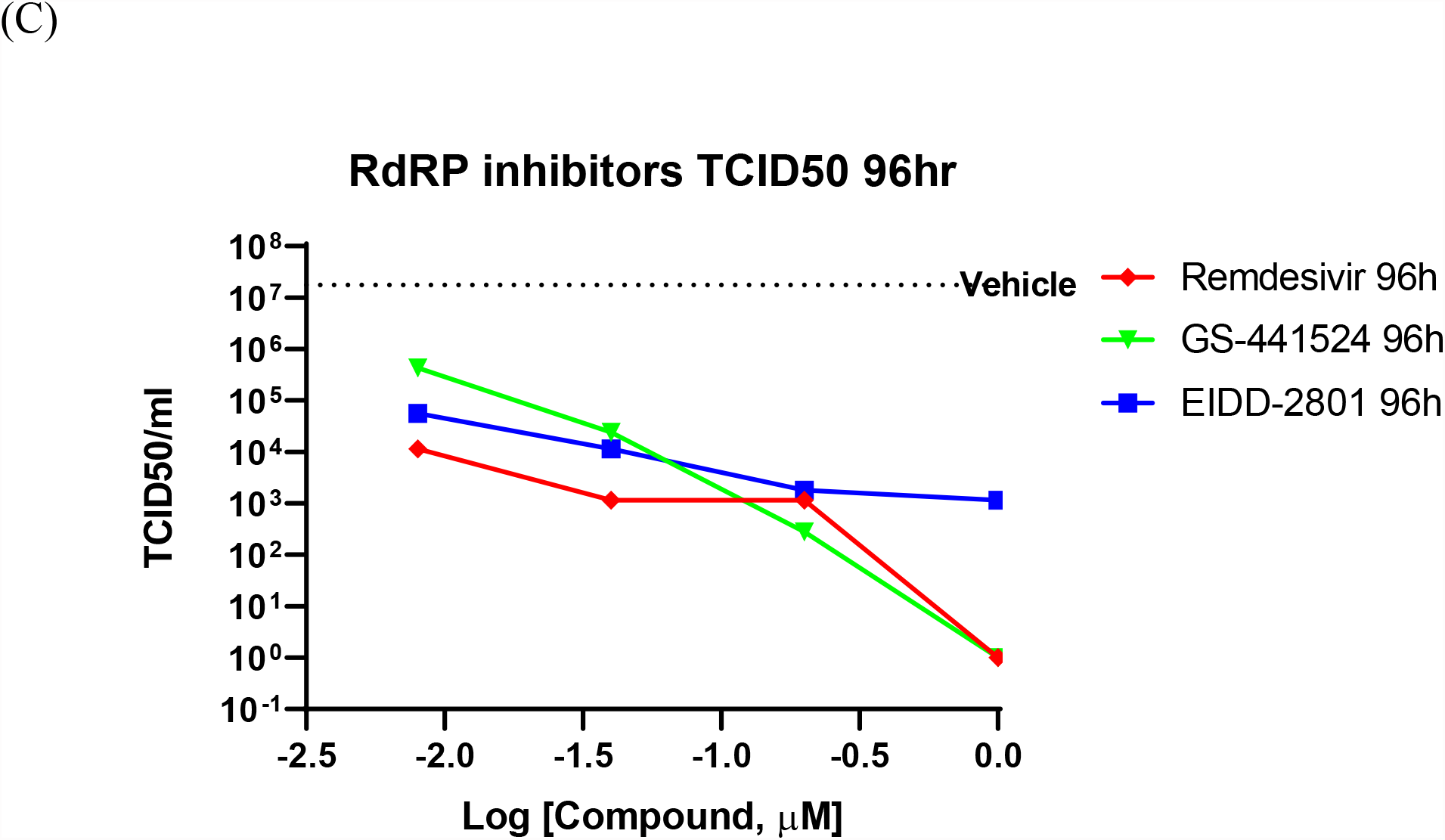
Anti-SARS-CoV-2 activity of GS-441524 and Remdesivir. (A) GS-441524 inhibited SARS-CoV-2 CPE in Vero E6 cells with an EC_50_ of 1.86 μM compared to remdesivir (EC_50_ of 7.43 μM). Neither compounds showed cytotoxicity at the concentrations used. (B) GS-441524 reduced viral load of SARS-CoV-2 infection in the 3D EpiAirway HAE tissue model. TCID50/mL was determined for remdesivir, GS-441524 and EIDD-2801 after (B) 24 h or (C) 96 h of SARS-CoV-2 infection.

### GS-441524 *in vitro* ADME properties

#### Kinetic solubility and PAMPA permeability

GS-441524 had low to moderate aqueous solubility of 35 - 52 µg/mL at pH 5 - 7.4 in Fasted and Fed State Simulated Intestinal Fluid (FaSSIF and FeSSIF) buffers, and high aqueous solubility >1 mg/mL at pH 1.6 in Simulated Gastric Fluid (SGF) buffers. A summary of the various solubility assays is provided in Table 1. GS-441524 also had a low apparent passive permeability value (P_app_<1 × 10^−6^ cm/sec) in the PAMPA assay.

**Table 1:** Summary of aqueous solubility for GS-441524. (Data presented are the average of duplicate samples)

| **Experiment Type** | **Matrix** | **Solubility in Aqueous Buffer (mg/mL)** |
| --- | --- | --- |
| Kinetic Solubility | pH 7.4 buffer | 0.042 |
| Equilibrium Solubility | pH 2 buffer | >1 |
|  | pH 5 buffer | 0.052 |
|  | pH 7.4 buffer | 0.047 |
|  | SGF | >1 |
|  | FaSSIF | 0.035 |
|  | FeSSIF | 0.052 |

#### Plasma protein binding and red blood cell (RBC)-to-plasma partitioning

Plasma protein binding studies and blood-to-plasma partitioning studies for GS-441524 were performed in human, rat, mouse, dog, and monkey (Table 2). Unbound fraction of GS-441524 was similar in rat, mouse, dog, and monkey (62-64%), whereas it was higher in human (∼78%). Overall, GS-441524 was a low-plasma protein bound drug candidate. Blood-to-plasma partitioning ratio for GS-441524 was ∼1.2 in all species tested (Table 2).

**Table 2:** Summary of *in vitro* ADME properties for GS-441524. (Data presented are the average of duplicate samples)

| **Experiment** | **Human** | **SD-Rat** | **CD-1**<br>**Mouse** | **Beagle**<br>**Dog** | **Cynomolgus**<br>**Monkey** |
| --- | --- | --- | --- | --- | --- |
| Microsomal Stability<br>$t_{1/2}$ (min) | >120 | 61±9 | 78±3 | >120 | >120 |
| Cytosol Stability<br>$t_{1/2}$ (min) | >120 | >120 | >120 | >120 | >120 |
| Hepatocyte Stability<br>$t_{1/2}$ (min) | >120 | >120 | >120 | >120 | >120 |
| Plasma Protein<br>Binding<br>(% bound) | 21.5 | 36.1 | 35.1 | 36.5 | 38.1 |
| Blood-to-Plasma<br>Partitioning (Ratio) | 1.18 | 1.18 | 1.13 | 1.20 | 1.18 |

#### Metabolic stability in liver microsomes, cytosols and hepatocytes

The susceptibility of GS-441524 to various drug metabolizing enzymes in the liver was evaluated using liver microsomal and cytosolic fractions, and hepatocytes in five species including human, rat, mouse, dog, and monkey. GS-441524 exhibited high stability in human, dog, and monkey microsomal fractions and moderate-high stability in rat and mouse microsomal fractions. GS-441524 was found to be stable in cytosolic fractions of all five species.

The stability of GS-441524 was also determined in human, rat, mouse, dog, and monkey hepatocytes (Table 2), and the depletion of GS-441524 was very low across five species for up to 2 h post incubation at 37°C, suggesting that GS-441524 is metabolically stable.

#### Metabolite identification in mouse, rat, dog, monkey, and human hepatocytes

GS-441524 was stable in mouse, rat, dog, monkey, and human plateable hepatocytes after incubation for 48 h. Testosterone was used as positive control for this assay and showed the expected results in all species, demonstrating that hepatocytes used in this study were metabolically active. There were no detectable phosphorylated metabolites in these cells under current experimental conditions; other potential metabolites, such as biotransformation via oxidation or oxidative deamination, were also not found in post-incubation samples. However, three possible glucuronidation metabolites and one possible sulfation metabolite were detected at trace amount based on mass spectrum analysis. Due to the low concentrations of these metabolites, it was not possible to observe them in LC-UV signal and to obtain reliable fragment ion information to propose possible structures of these metabolites.

#### Transporter-mediated transport of GS-441524

For efflux transporter assessment, bidirectional transport assays were conducted using Caco-2, MDCK-MDR1 and MDCK-BCRP monolayer cell models. The efflux ratios (P_app A to B_/ P_app B to A_) with or without transporter inhibitors were determined. All the monolayer cell models were validated using digoxin and cladribine as the positive controls of MDR1 and BCRP substrates, respectively. The efflux ratios of digoxin were 27 and 110 in Caco-2 and MDCK-MDR1 cells, while those of cladribine were 26 and 153 in Caco-2 and MDCK-BCRP cells, respectively. Recovery of GS-441524 was in the acceptable range (75 - 125%) in all efflux transporter assays.

In Caco-2 cells, the permeability from apical to basolateral side (P_app A to B_) of GS-441524 was 0.099×10^−6^ cm/s. The efflux ratio of GS-441524 without any inhibitor was 9.78 in Caco-2 cells, and the addition of the MDR1 inhibitor (1 µM Valspodar) reduced the efflux ratio to 4.93. In MDCK-MDR1 cells, efflux of GS-441524 was completely inhibited by pre-incubation with Valspodar (Table 3), suggesting GS-441524 is a substrate for the MDR1 transporter.

**Table 3.**
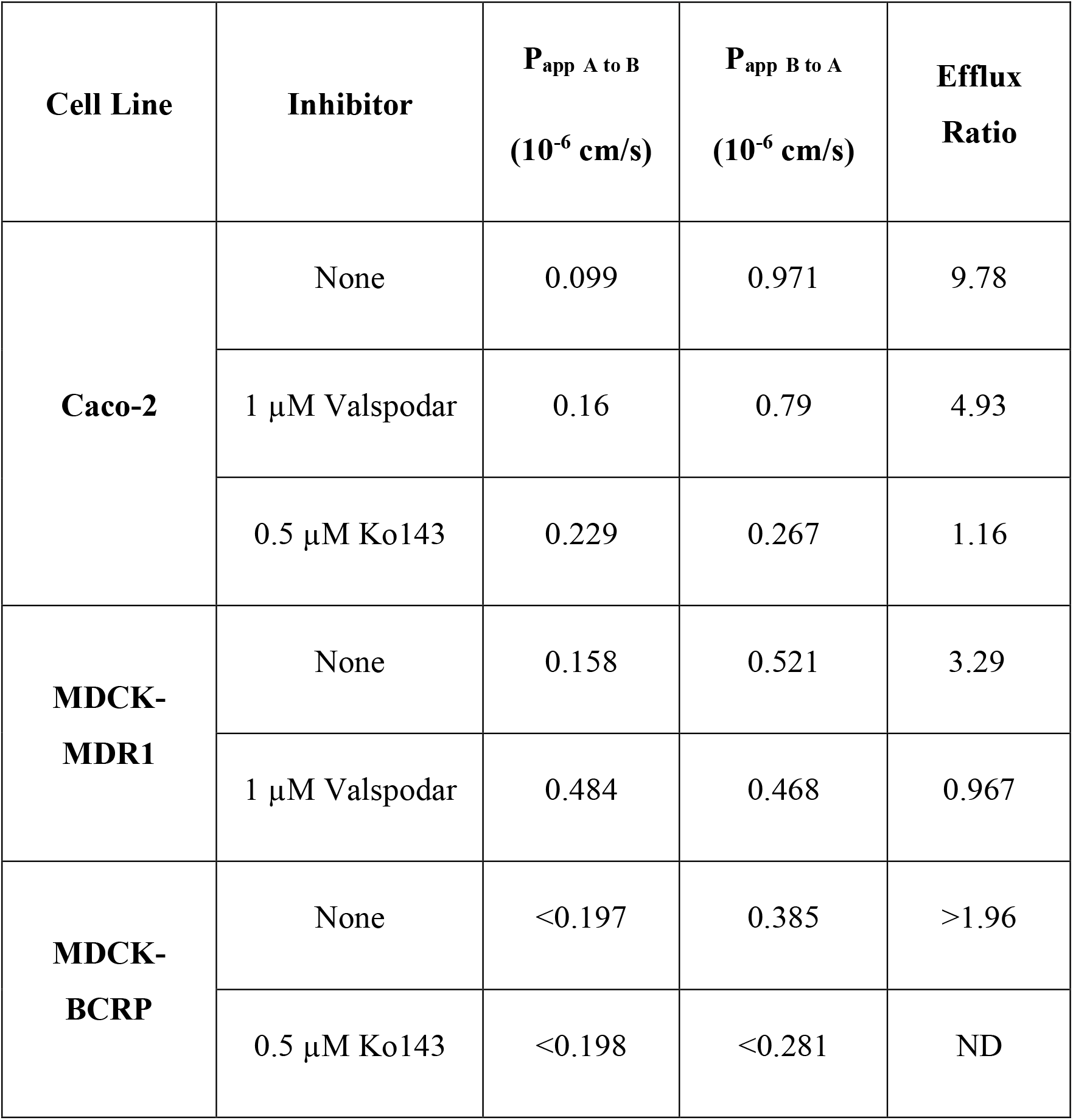
Active efflux of GS-441524 (5 µM) in Caco-2, MDCK-MDR1 and MDCK-BCRP cell monolayers with the present and absence of Valspodar (MDR1 inhibitor) and Ko143 (BCRP inhibitor) (Data presented are the average of duplicate samples)

| **Cell Line** | **Inhibitor** | **P<sub>app</sub> A to B**<br>**(10<sup>-6</sup> cm/s)** | **P<sub>app</sub> B to A**<br>**(10<sup>-6</sup> cm/s)** | **Efflux**<br>**Ratio** |
| --- | --- | --- | --- | --- |
| **Caco-2** | None | 0.099 | 0.971 | 9.78 |
| | 1 μM Valspodar | 0.16 | 0.79 | 4.93 |
| | 0.5 μM Ko143 | 0.229 | 0.267 | 1.16 |
| **MDCK-MDR1** | None | 0.158 | 0.521 | 3.29 |
| | 1 μM Valspodar | 0.484 | 0.468 | 0.967 |
| **MDCK-BCRP** | None | <0.197 | 0.385 | >1.96 |
| | 0.5 μM Ko143 | <0.198 | <0.281 | ND |

We further investigated the potential roles of BCRP on the transport of GS-441524. Pre-incubation of BCRP inhibitor (0.5 µM Ko143) in Caco-2 cells completely reduced the efflux ratio from 9.78 to 1.16 (Table 3). In MDCK-BCRP cells, the permeability from apical to basal side (P_app A to B_) for GS-441524 was too low to be determined; thus, the efflux ratio could not be calculated. The permeability from basal to apical side (P_app B to A_) decreased after pre-incubation of Ko143 (Table 3), suggesting GS-441524 could be a substrate of BCRP.

GS-441524 was further screened for six major uptake human nucleoside transporters, namely ENT1, ENT2, ENT4, CNT1, CNT2, and CNT3, on MDCK cells at concentrations of 10, 30 and 100 µM (Fig. 2). GS-441524 is a substrate for CNT3 and ENT1. Compared to matched control cells, the uptake of GS-441524 was 81.2 to 164-fold and 3.38 to7.45-fold higher in CNT3- and ENT1-transfected cells, respectively. Upon addition of the reference inhibitor adenosine, 98.9% and 92.3% inhibition of this uptake was seen in CNT3- and ENT1-transfected cells, respectively. This confirms that GS-441524 is a substrate for CNT3 and ENT1.

**Figure 2.**
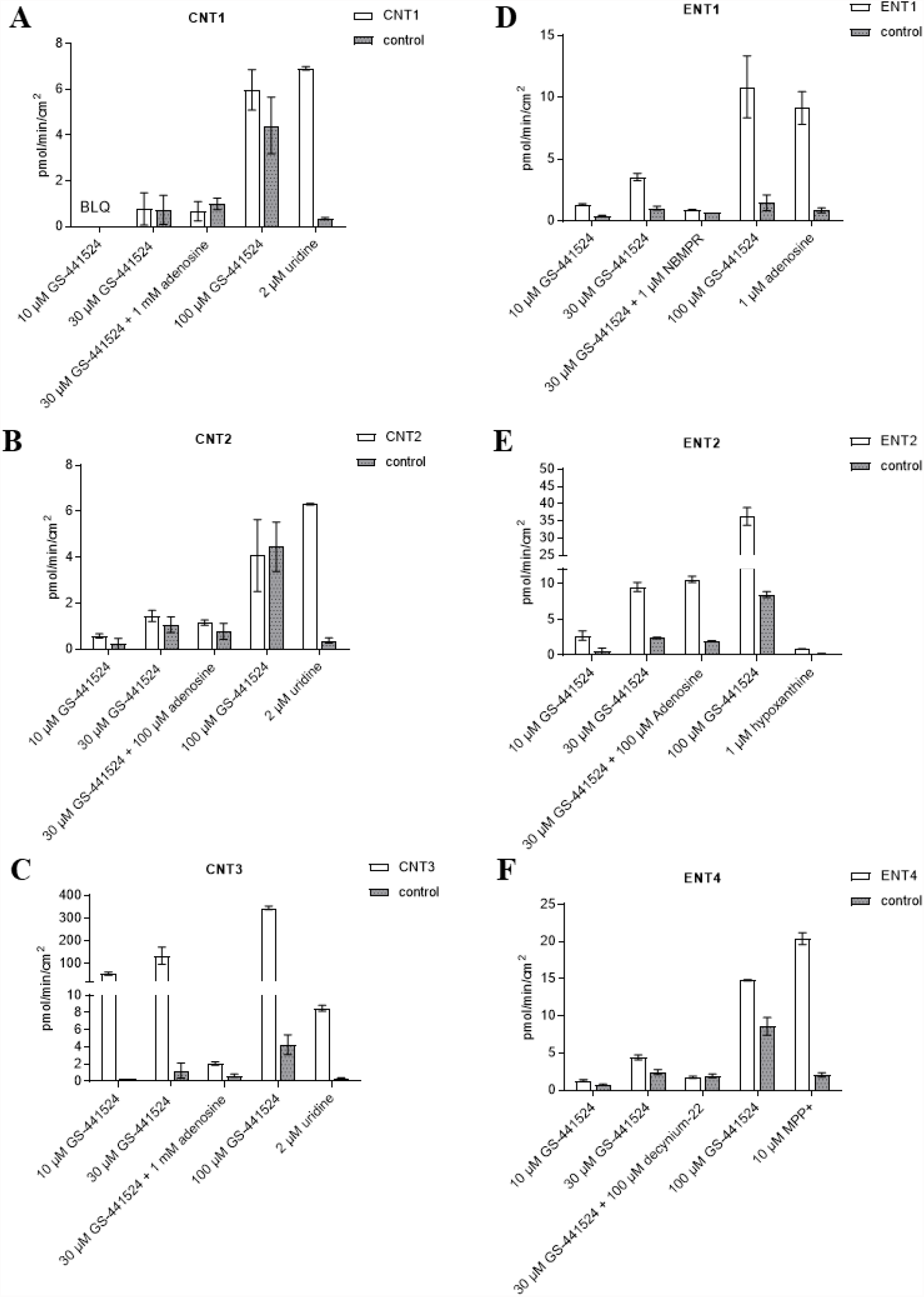
Human nucleoside transporter uptake of GS-441524. (A) CNT1, (B) CNT2, (C) CNT3, (D) ENT1, (E) ENT2, and (F) ENT4. (Data represent the mean and standard deviation of triplicate samples. BLQ: Below limit of quantitation, LOQ= 0.222 pmol/min/cm^2^)

GS-441524 is unlikely to be a substrate for CNT1, ENT4, and CNT2 under the study conditions because less than a 2-fold [20] difference in uptake for CNT1 and ENT4 was observed in transporter-transfected cells compared to control cells. In CNT2-transfected cells, uptake of GS-441524 was only slightly higher than 2-fold at 10 µM and was less than 2-fold at 30 and 100 µM.

In ENT2-transfected cells, uptake of GS-441524 was 3.95- to 4.7-fold higher compared to that in the matched control cells, suggesting that GS-441524 is a substrate for ENT2. However, upon addition of the reference inhibitor adenosine, no inhibition was observed, indicating that ENT2-mediated uptake of GS-441524 may not be sensitive to inhibition by 100 µM adenosine.

### UPL-MS/MS bioanalysis method for GS-441524

An accurate, rapid, and selective UPLC-MS/MS method was developed to quantify GS-441524 in plasma, urine, and bile samples. GS-441524 had poor retention on regular C_18_ columns due to its hydrophilicity. Several UPLC columns were evaluated, including HILIC columns. Good chromatographic retention was achieved on an Atlantis™ PREMIER BEH C_18_AX column (2.1 × 50 mm). The retention time for GS-441524 and SIL-IS (internal standard, ^13^C_5_-GS-441524) was 0.9 min with the gradient elution at the flow rate of 0.5 mL/min. For mass spectrometric detection, selected reaction monitoring (SRM) was used in the positive ionization mode. No endogenous components interfered with the analysis of GS-441524 and the internal standard in PK samples. The correlation coefficient of the calibration curves was greater than 0.99.

### Pharmacokinetics of GS-441524 in mice, rats, monkeys, and dogs

In C57BL/6 mice, GS-441524 was administered at 5 mg/kg IV and 10 mg/kg PO to evaluate its bioavailability and PK profiles. Following IV administration, the clearance of GS-441524 determined with plasma drug concentrations was 26 mL/min/kg and the steady state volume of distribution (Vd_ss_) was 2.4 L/kg. For 10 mg/kg PO administration, the peak plasma concentration was achieved at 1.5 h (T_max_ = 1.5 h; Fig. 3) with the maximum concentration (C_max_) of 582 ng/mL. The half-life (t_1/2_) was 3.9 h and the bioavailability was 39% with area under the curve (AUC) of 2540 ng·h/mL. The results are summarized in Table 4.

**Table 4.**
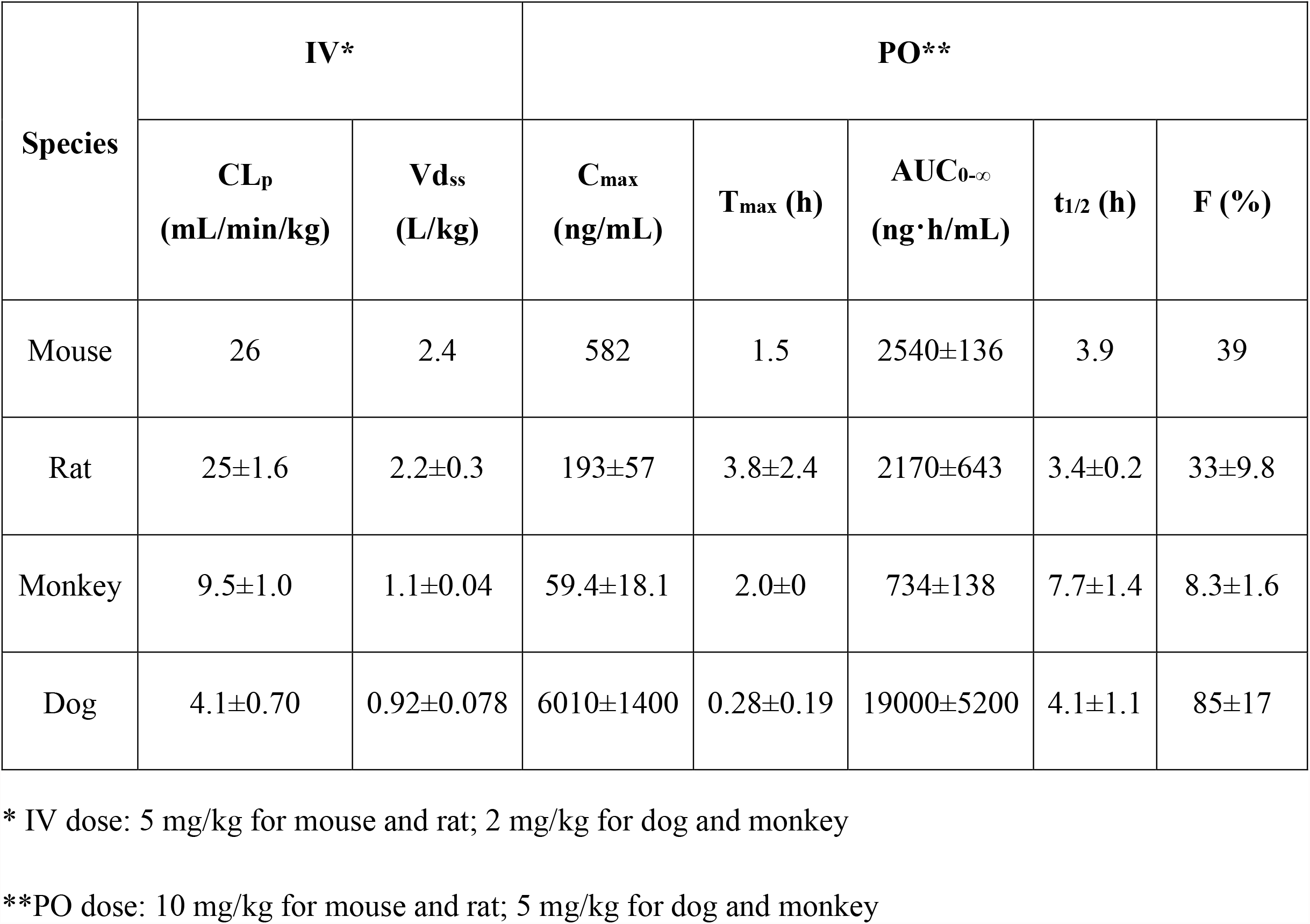
Pharmacokinetic parameters of GS-441524 in mice, rats, monkeys, and dogs after single IV and PO administration.

| Species | IV* |  | PO** |  |  |  |  |
| --- | --- | --- | --- | --- | --- | --- | --- |
|  | CL<sub>p</sub><br>(mL/min/kg) | Vd<sub>ss</sub><br>(L/kg) | C<sub>max</sub><br>(ng/mL) | T<sub>max</sub> (h) | AUC<sub>0-∞</sub><br>(ng·h/mL) | t<sub>1/2</sub> (h) | F (%) |
| Mouse | 26 | 2.4 | 582 | 1.5 | 2540±136 | 3.9 | 39 |
| Rat | 25±1.6 | 2.2±0.3 | 193±57 | 3.8±2.4 | 2170±643 | 3.4±0.2 | 33±9.8 |
| Monkey | 9.5±1.0 | 1.1±0.04 | 59.4±18.1 | 2.0±0 | 734±138 | 7.7±1.4 | 8.3±1.6 |
| Dog | 4.1±0.70 | 0.92±0.078 | 6010±1400 | 0.28±0.19 | 19000±5200 | 4.1±1.1 | 85±17 |

\* IV dose: 5 mg/kg for mouse and rat; 2 mg/kg for dog and monkey
\*\*PO dose: 10 mg/kg for mouse and rat; 5 mg/kg for dog and monkey

**Figure 3.**
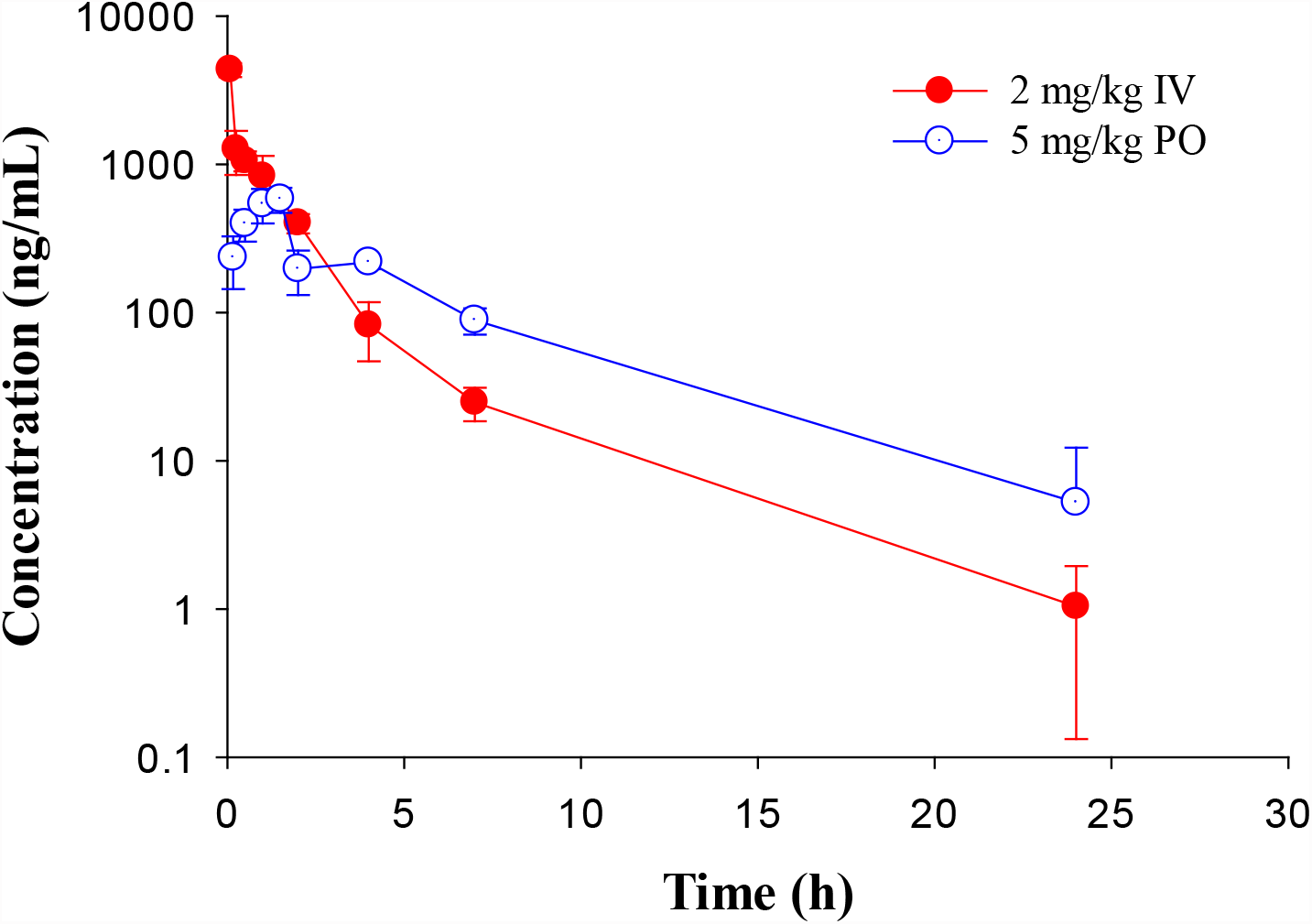
Pharmacokinetic profiles of GS-441524 in C57BL/6 mice. Plasma concentrations of GS-441524 following intravenous (IV) administration of 5 mg/kg and oral (PO) administration of 10 mg/kg. Data are the mean ± SD (n=3/time point).

In Sprague Dawley rats, GS-441524 was administered at 1 and 5 mg/kg IV; and 10, 30 and 100 mg/kg PO to evaluate its PK profiles (Fig. 4). PK parameters are summarized in Table 4. For IV administrations of 1 and 5 mg/kg, the clearances were 31 and 25 mL/min/kg, and Vd_ss_ were 2.6 and 2.2 L/kg, respectively (Table 5). After oral administration of 10 mg/kg, the bioavailability was estimated to be 33%. The C_max_ and AUC increased with the dose in a less than dose-proportional manner after 10, 30 and 100 mg/kg PO administration. The half-life (t_1/2_) was 3.4 - 4.9 h. Following IV administration of 2 mg/kg in bile duct cannulated (BDC) rats, urinary and biliary excretions of unchanged parent drug were 65% and 0.8% of dose, respectively, suggesting that urinary excretion is a major elimination route for GS-441524 in rats.

**Table 5.** Pharmacokinetic parameters of GS-441524 in rats after single PO administration. (Mean ± SD)

| **Dose**<br>**(mg/kg)** | **C<sub>max</sub>**<br>**(ng/mL)** | **T<sub>max</sub> (h)** | **AUC<sub>0-∞</sub>**<br>**(ng·h/mL)** | **t<sub>1/2</sub> (h)** | **F (%)** |
| --- | --- | --- | --- | --- | --- |
| 10 | 193±57 | 3.8±2.4 | 2170±643 | 3.4±0.2 | 33±9.8 |
| 30 | 326±77 | 3.8±2.4 | 4220±746 | 3.4±0.2 | 21±3.8 |
| 100 | 825±91 | 2.3±1.3 | 9560±2130 | 4.2±1.8 | 15±3.2 |

**Figure 4.**
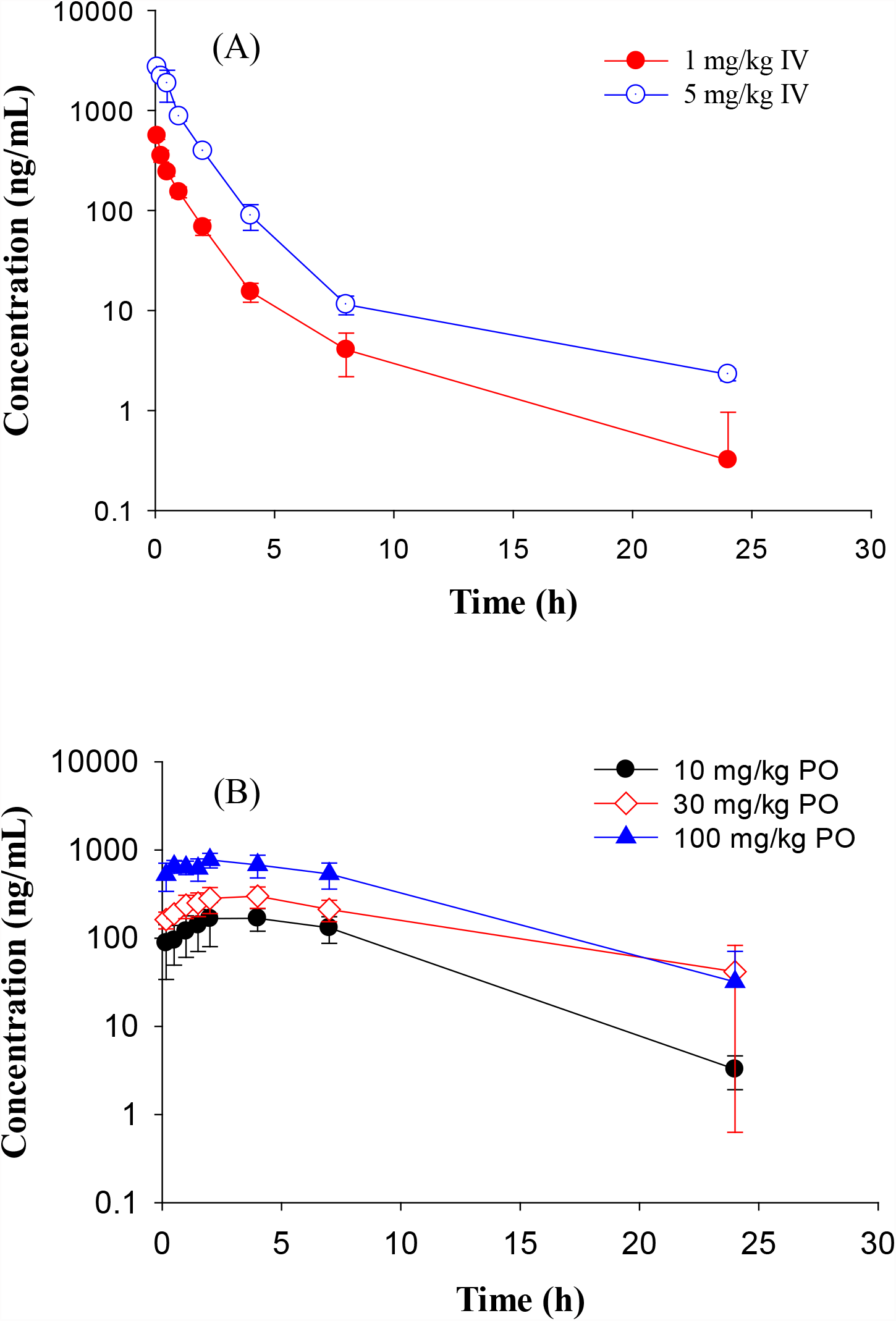
Pharmacokinetic profiles of GS-441524 in Sprague Dawley rats. Plasma concentrations of GS-441524 following a single intravenous (IV) of 1 and 5 mg/kg (A) or oral (PO) dose of 10, 30 and 100 mg/kg (B). Data are the mean ± SD (n=3-4/group).

In Cynomolgus monkeys, a single dose of GS-441524 (2 mg/kg IV and 5 mg/kg PO) was administrated (Fig. 5). The mean clearance from three animals was 9.5 mL/min/kg and the mean steady state volume of distribution was 1.1 L/kg. C_max_, AUC and t_1/2_ are summarized in Table 4. Following PO administration, the bioavailability was 8.3%, the lowest in the four animal species evaluated. Following IV administration of 2 mg/kg, the mean urinary excretion of unchanged parent drug was 17% over a collection period of 48 h.

**Figure 5.**
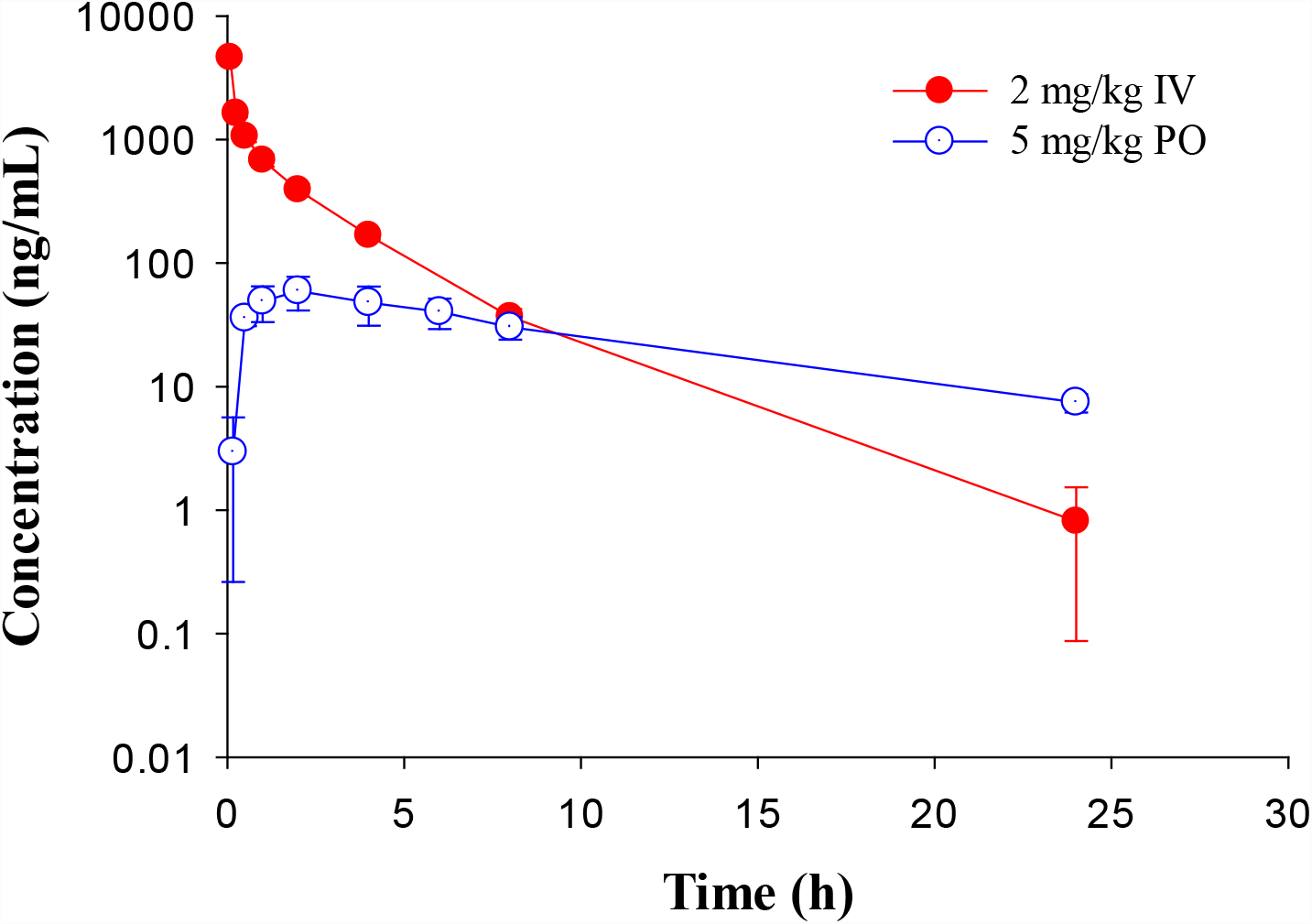
Pharmacokinetic profiles of GS-441524 in Cynomolgus monkeys. Plasma concentrations of GS-441524 following intravenous (IV) administration of 2 mg/kg and oral (PO) administration of 5 mg/kg. Data are the mean ± SD (n=3/group).

In Beagle dogs, a single dose of GS-441524 (2 mg/kg IV and 5 mg/kg PO) was administrated (Fig. 6). The mean clearance from three animals was 4.1 mL/min/kg and the mean steady state volume of distribution was 0.92 L/kg. Following PO administration, the bioavailability was 85%, the highest in four animal species evaluated. The t_1/2_ value was ∼ 4 h for both IV and PO administration (Table 4). Following IV administration of 2 mg/kg, the mean urinary excretion of unchanged parent drug was 64%, suggesting that urinary excretion is a major elimination route for GS-441524 in dogs.

**Figure 6.**
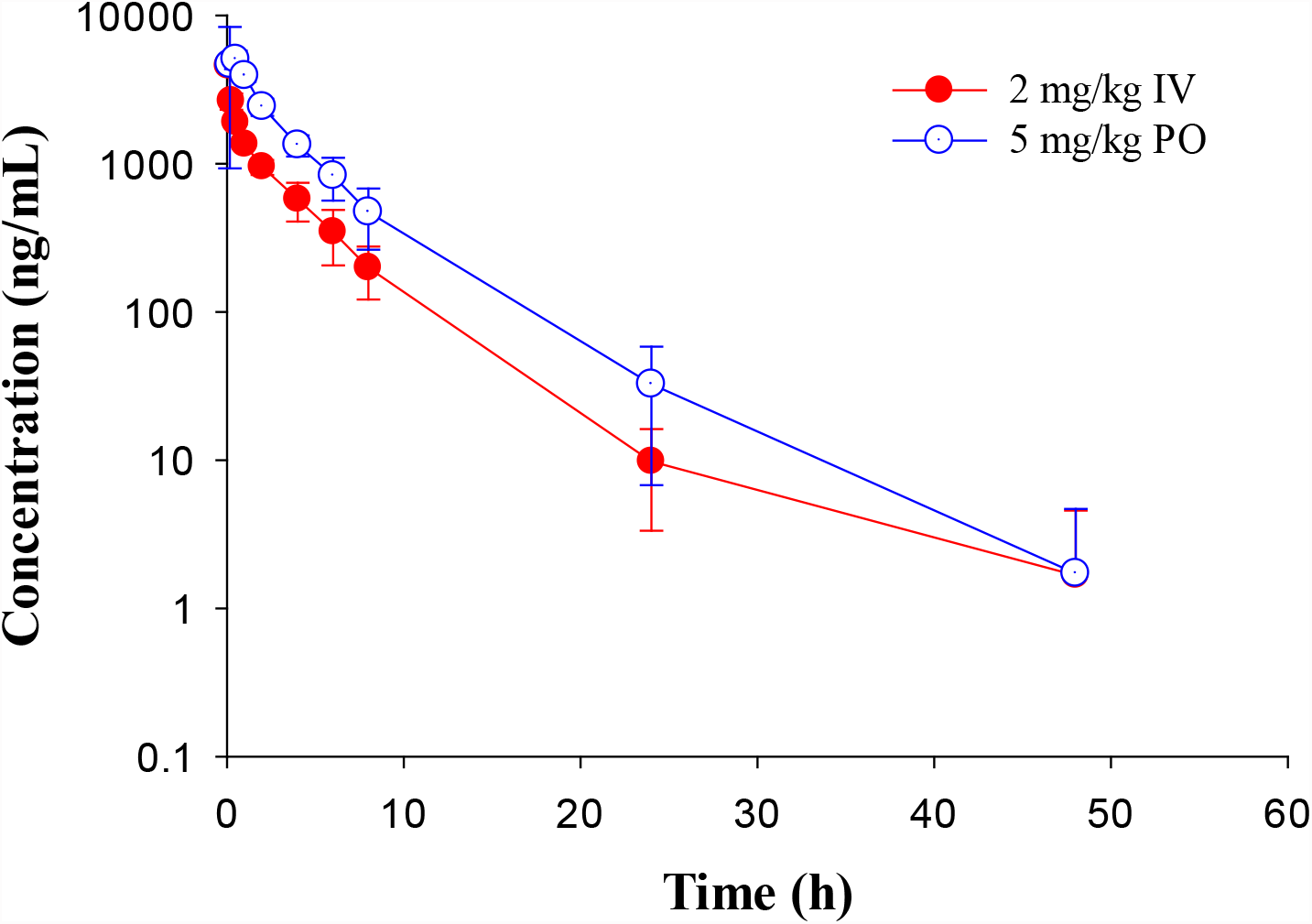
Pharmacokinetic profiles of GS-441524 in Beagle dogs. Plasma concentrations of GS-441524 following intravenous (IV) administration of 2 mg/kg and oral (PO) administration of 5 mg/kg. Data are the mean ± SD (n=3/group).

### GS-441524 human PK prediction

To predict human exposures after oral administration, human PK parameters were estimated using allometric scaling and human PK profile was simulated using GastroPlus. CL_p_ and Vd_ss_ from mouse, rat, monkey, and dog parameters summarized in Table 4 were used. The allometric scaling was performed as previously described [21]. From the linear regression of log-log curves of CL_p_ and Vd_ss_ vs body weight of 4 species (Fig. 7), we derived the coefficient (a) and exponent (b) of the allometric equation Y = a·W^b^ where Y is CL_p_ or Vd_ss_, and W is body weight [21]. The derived allometric scaling equations were defined as CL_p_(mL/min) = 12.6·BW^0.64^ and Vd_ss_ (L) = 1.5·BW^0.8^. Using the allometric scaling equation, the predicted human CL_p_ is 2.7 mL/min/kg and Vd_ss_ is 0.6 L/kg, and these values were used for human PK simulation.

**Figure 7:**
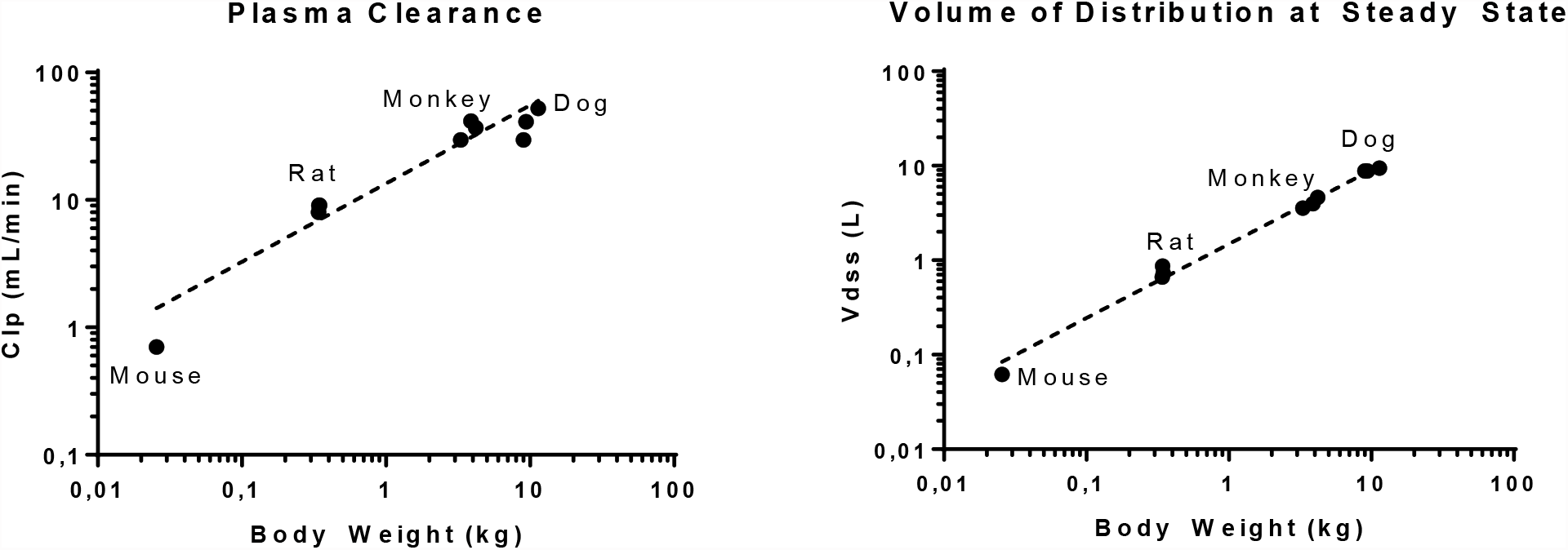
Log-log plot of plasma clearance and volume of distribution at steady state of mouse, rat, monkey, and dog *vs* respective body weights. Data presented as individual value.

To simulate the PK profile in humans, we used a simple one compartment model in conjunction with the advanced compartmental absorption and transit model in GastroPlus. The experimental parameters used in these predictions include plasma protein binding, RBC to plasma partitioning, equilibrium solubility at pH 7.4 and Caco-2 P_app A to B_ permeability. Additionally, the clearance and volume of distribution obtained from allometric scaling were used as inputs for these predictions. Since there was a large variation in bioavailability amongst preclinical species (8% - 85%), we capped bioavailability at 20% for our prediction as a conservative approach. Different dosing regimens were used to achieve plasma concentrations above 0.5 µM, 1 µM and 2 µM. Our predictions reveal that 300 mg, 600 mg, and 1000 mg b.i.d. dosing would be able to achieve plasma concentrations of 0.5 µM, 1 µM, and 2 µM respectively, throughout the dosing regimen (Fig. 8). A summary of PK parameters obtained from the predictions are presented in Table 6.

**Table 6.** Human pharmacokinetic parameters of GS-441524 predicted using GastroPlus.

| Dosing Regimen<br>(mg, b.i.d) | Day One |  |  |  | Steady State |  |  |  |
| --- | --- | --- | --- | --- | --- | --- | --- | --- |
|  | C<sub>max</sub> | C<sub>trough</sub> | Daily AUC* | C<sub>mean</sub> ** | C<sub>max</sub> | C<sub>trough</sub> | Daily AUC* | C<sub>mean</sub> ** |
|  | (μM) | (μM) | (μM·h) | (μM) | (μM) | (μM) | (μM·h) | (μM) |
| 300, b.i.d | 2.1 | 0.4 | 29 | 1.2 | 2.2 | 0.4 | 30 | 1.3 |
| 600, b.i.d | 4.0 | 0.8 | 60 | 2.5 | 4.2 | 0.8 | 63 | 2.6 |
| 1000, b.i.d | 7.2 | 1.6 | 112 | 4.7 | 7.6 | 1.7 | 118 | 4.9 |

\*Daily AUC: AUC of two doses
\*\*C<sub>mean</sub>: Daily AUC/24 h

**Figure 8:**
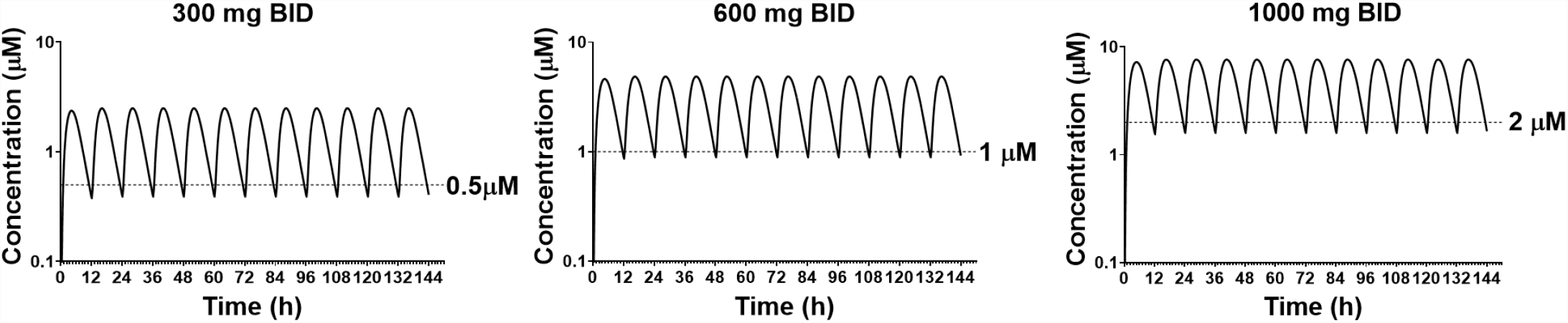
Predicted plasma concentrations of GS-441524 using single compartment model with advanced compartmental absorption and transit module in GastroPlus.

## DISCUSSION

Remdesivir has shown to reduce progression to hospitalization or death by 87% for the treatment of high risk nonhospitalized COVID-19 patients [3]. However, remdesivir can only be given by IV administration [4] which significantly limits its use [5]. With the aim of assessing whether GS-441524 could be potentially used for oral administration, in this study, we determined the *in vitro* potency, *in vitro* ADME properties, and preclinical PK of GS-441524. The *in vitro* potency results confirmed that activity of GS-441524 against SARS-CoV-2 was slightly more potent in Vero E6 cells but similar to that of remdesivir in the 3D HAE tissue model [22, 23]. Furthermore, the preclinical PK evaluations indicated that GS-441524 is orally bioavailable.

Like other antiviral nucleoside analogs, GS-441524 demonstrated very limited membrane permeability. Therefore, active transport may play a significant role in the absorption and disposition of GS-441524. As a major metabolite of remdesivir, GS-441524 was not a substrate of the following human transporters: OCT1, OCT2, OAT1, OAT3, OATP1B1, OATP1B3, MATE1 and MATE2-k [26]. The roles of MDR1, BCRP, concentrative nucleoside transporters (CNTs), and equilibrative nucleoside transporters (ENTs) for GS-441524 were unknown. Our *in vitro* results demonstrated that GS-441524 was a substrate of MDR1, BCRP, CNT3, ENT1 and ENT2, but not a substrate of CNT1, CNT2, and ENT4. The expression of ENT1 and ENT2 was comparable to that of MDR1 and BCRP in human kidney tissue [28, 34], and the uptake of GS-441524 by ENT1 and ENT2 could offset part of the efflux by MDR1 and BCRP. Since CNT3’s activity in the uptake of GS-441524 was 164 times that of control cells, CNT3 may play a substantial role in the reabsorption of GS-441524 in kidney. In addition, both MDR1 and BCRP are expressed in the apical side of cells in liver and brain [20, 28], and CNT3, ENT1, and ENT2 are ubiquitously expressed in various tissues [28, 34]. These transporters may also play a role in the distribution of GS-441524. ENT1 is also expressed on the mitochondria membrane and has been linked to the mitochondria toxicity of some antiviral drugs [43]. In our 7-day dose range finding studies in rats, dogs, and monkeys, orally administered GS-441524 was well tolerated at the maximum feasible dose of 1500, 2000 and 1000 mg/kg/day, respectively (https://opendata.ncats.nih.gov/covid19/GS-441524).

PK studies of GS-441524 were performed to investigate the absorption, distribution, clearance, and *in vivo* exposure after IV and PO administration in mice, rats, monkeys, and dogs. The plasma clearance values ranged from 4.1 mL/min/kg in dogs to 26 mL/min/kg in mice, which were considered as low to moderate clearance when compared to the hepatic blood flow in these species, after correcting for the blood/plasma ratio. The oral bioavailability had large species differences, ranging from 8.3% in monkeys to 85% in dogs (Table 4), likely caused by the variability in animal physiology (e.g., GI transit time, pH, etc.) and drug physicochemical properties. MDR1, BCRP, ENT1, and ENT2 showed large species difference in their endogenous or exogenous substrate profiles or in their substrate affinity [30-32, 35-37].

The AUC of GS-441524 increased with the dose in a less than dose-proportional manner after 10, 30 and 100 mg/kg PO administration in rats (Table 5), likely due to the solubility limit and the saturated CNT3 activity. Following a single 100 mg/kg oral dose of GS-441524 in rats, the systemic exposure (AUC) was 9560 ng·h/mL, higher than the AUC of GS-441524 (7350 ng·h/mL) following a 225 mg IV remdesivir dose in human [44].

The urinary excretion of GS-441524 was the predominant elimination route in rats, as ∼ 65% of GS-441524 was recovered in urine and <1% was recovered in bile over a collection period of 48 h after IV administration in BDC rats. Similarly, ∼64% of GS-441524 was recovered in dog urine. However, only 17% of GS-441524 was recovered in monkey urine over a collection period of 48 h after IV administration. The low urinary excretion in monkey might be related to species difference in the transporters’ affinity to GS-441524.

The human CL_p_ and Vd_ss_ were calculated using allometric scaling of the animal CL_p_ and Vd_ss_ from the PK studies. The calculated human CL_p_ is low, approximately 10% of the human hepatic blood flow. Additionally, the human PK profiles of GS-441524 was simulated from the experimental *in vitro* data and the calculated CL_p_ and Vd_ss_ values using GastroPlus software. The prediction indicates that 1000 mg, b.i.d. oral dosing regimen would be sufficient to achieve plasma concentrations above the EC_50_ (1.86 μM) in the SARS-CoV-2 live virus assay.

Clinically, antiviral treatment usually involves more than one therapeutic. Since GS-441524 does not undergo extensive metabolism, transporters may be an important factor mediating related drug-drug interactions. Inhibition of MDR1 and BCRP has significant impact on the clinical PK of co-administered antiviral agents in some antiviral regimens [38], and inhibition of ENT1 and ENT2 leads to accumulation of nucleoside analogs inside the human small intestine [39]. Since molnupiravir and its active metabolite EIDD-1931 have been reported as inhibitors of ENT1 and ENT2 [27], drug-drug interaction potential needs to be assessed if a combination therapy of molnpiravir and redemsivir/GS-441524 is applied. Under our *in vitro* study conditions, 100 μM adenosine did not inhibit the uptake of GS-441524 by ENT2, and 1 mM adenosine inhibited the uptake by CNT3. Thus, inflammation-related fluctuation of plasma adenosine level (10-to 20-fold higher than the normal concentration at 0.4 – 0.8 μM) [40] is unlikely to alter the cross-membrane transport of GS-441524.

## CONCLUSIONS

We demonstrated that GS-441524, the active metabolite of remdesivir, could be an oral drug candidate as an antiviral agent for SARS-CoV-2. GS-441524 had a low to moderate plasma clearance in mouse, rat, cynomolgus monkeys, and dogs. The free fractions of GS-441524 were 62-78% in plasma protein binding assay in all studied species. Renal excretion plays an important role in the elimination of GS-441524. The *in vitro* ADME and PK results from this study allowed us to propose human PO dosing regimens compatible with b.i.d. dose scheduling. A simulation of b.i.d dose of 1000 mg would achieve plasma concentration of 2 μM, above EC_50_ in the Vero E6 and 3D HAE tissue model.

## MATERIALS AND METHODS

Formic acid and LC-MS grade acetonitrile were obtained from Fisher Scientific (Pittsburgh, PA, U.S.A.). Isopropyl alcohol and LC-MS grade methanol and acetonitrile were obtained from Sigma-Aldrich (St. Louis, MO, U.S.A.). Deionized water was generated using a Milli-Q Ultrapure water purification system from EMD Millipore (Billerica, MA, U.S.A.). GS-442524 and ^13^C_5_-GS-441524 (SIL-internal standard, SIL-IS) were ordered from AK Scientific (Union City, CA) and Cambridge Isotope Laboratories, Inc. (Tewksbury, MA), respectively. Control male C57BL6 mouse, Sprague Dawley rat, Cynomolgus monkey and Beagle dog plasmas were ordered from Bioreclamation IVT (Westbury, NY, U.S.A.).

### SARS-CoV-2 infection of Vero E6 cells measured by cytopathic effect (CPE)

The CPE assay was performed by the contract BSL-3 laboratory (Southern Research Institute) as described previously [19, 45]. Briefly, Vero E6 cells previously selected for high ACE2 expression were cultured in MEM plus 2% heat-inactivated FBS supplemented with 1% Pen/Strep, 1% GlutaMax, and 1% HEPES. The cells were inoculated at a MOI of 0.002 with SARS-CoV-2 (USA_WA1/2020) in media and quickly dispensed into 384-well assay plates containing test compounds at 30 μL/well (4,000 cells/well). After 72 h incubation at 37°C, 5% CO_2_, and 90% humidity, 30 μL/well of the CellTiter-Glo ATP content assay reagent mixture (Promega, #G7573) was added that was incubated for 10 min at room temperature followed by a detection of luminescence signal on a Perkin Elmer Envision or BMG CLARIOstar plate reader. The compound cytotoxicity was detected using the same protocol as CPE assay in the absence of SARS-CoV-2 virus. The assay ready plate containing compounds was prepared using an Echo acoustic dispensing instrument (Labcyte) that had 90 nL aliquot of each diluted sample per well. The final DMSO concentration in the assay was 0.3%.

### SARS-CoV-2 infection in a 3D EpiAirway model

The 3D EpiAirway infection model was contracted to the Center for Predictive Medicine for Biodefense and Emerging Infectious Diseases at the Louisville University. Differentiated human tracheobronchial epithelial tissues (EpiAirway™ AIR-100) were obtained from MatTek Corporation (Ashland, MA). EpiAirway tissues were grown on 6 mm^2^ mesh disks in transwell inserts. Three days prior to shipment, the tissues were transferred into hydrocortisone-free medium. The tissues were shipped on a sheet of agarose, which was removed upon receipt, and the tissues were cultured on inserts at ALI. One insert is estimated to consist of approximately 1.2 × 10^6^ cells. Tissues were pretreated with compounds for 1 h on the apical and basal sides. After compound pretreatment, liquid was removed from the apical side, and virus (2019-nCoV/USA-WAl/2020 at MOI of 0.1) was inoculated in 0.15 mL assay medium onto the apical layer for 1 ho. Following 1 h inoculation, virus-containing media was removed from apical layer and the apical side was washed with 400 μl TEER buffer. Media on the basolateral side was replaced with fresh compound-containing maintenance medium at 24 h, 48 h and 72 h post-infection. At 24 h and 96 h post-infection, samples from the apical layer of the tissue were collected by washing layer with 400 μl TEER buffer and aliquoted for TCID_50_ (50% Tissue Culture Infectious Dose) determination. Media from the basal layer of the tissue were collected at the same time points for LDH cytotoxicity measurements (LDH-Glo Cytotoxicity Assay, Promega). TCID_50_ assay was performed as previously described [19]. The apical layer supernatant samples were diluted at 1 to 8 ratio that were added to 96-well assay plates containing 10,000/well Vero E6 cells, followed by incubation at 37°C, 5% CO_2_ and 95% relative humidity. After an incubation for 3 days (72 ± 4 h), crystal violet staining was performed to measure cytopathic effect (CPE) in the assay plates and the virus titers were calculated using the method of Reed and Muench [46]. Each of TCID_50_ values were determined in duplicates.

### Evaluation of *In vitro* ADME properties of GS-441524

#### Aqueous solubility

Kinetic aqueous solubility was determined at NCATS using Pion’s patented µSOL assay as described previously [47].

Equilibrium solubility was measured in pH 2.0, 5.0, 7.4 aqueous buffers as well as Simulated Gastric Fluid (SGF; pH 1.6), Fasted State Simulated Intestinal Fluid (FaSSIF; pH 6.5) and Fed State Simulated Intestinal Fluid (FeSSIF; pH 5.8). Briefly, >1 mg of GS-441524 powder was combined with 1 mL of buffer to make a >1 mg/mL mixture. Samples were shaken on a Thermomixer® overnight at room temperature and then passed through a 0.45 μm PTFE syringe filter. The filtrate was then sampled and diluted in a 1:1 mixture of buffer:ACN prior to analysis by UPLC-MS/MS.

#### Parallel artificial membrane permeability assay (PAMPA)

Stirring double-sink PAMPA (patented by pION Inc.) was employed to determine permeability. Assay was performed as described previously [48].

#### Metabolic stability assays

In the microsomal stability experiment, each reaction mixture (110 μL) consisted of GS-441524 (1 μM), human/S.D rat/CD-1 mouse/Gottingen minipig/Cynomolgus monkey/Beagle dog (Sekisui XenoTech, LLC) microsomal fractions (0.5 mg/mL), and NADPH regenerating system (1 μM) in phosphate buffer at pH 7.4. In the Cytosol stability experiment, each reaction mixture (110 μL) consisted of GS-441524 (1 μM) and human/S.D. rat/CD-1 mouse/Gottingen minipig/Cynomolgus monkey/Beagle dog (Sekisui XenoTech, LLC) cytosol fractions (2 mg/mL) in phosphate buffer (100 mM) at pH 7.4. Reaction mixtures were incubated in 384-well plates at 37°C for 0, 5, 10, 15, 30 and 60-min. The reaction was terminated by adding three volumes of cold ACN with internal standard (IS). Sample were analyzed using UPLC-MS/MS. Analysis and half-life calculations were performed using a previously described method [49, 50]

Cryopreserved mixed-gender human, male Sprague-Dawley rat, male CD-1 mouse, male Beagle dog, and male Cynomolgus monkey hepatocytes (Sekisui XenoTech, LLC) were thawed and equilibrated at 37°C. The reaction was initiated by spiking the test compound into the hepatocyte suspension at a 1 µM final GS-441524 concentration. The reaction mixture was incubated in a shaking water bath at 37°C and aliquots were withdrawn at 0, 5, 15, 30, 60 and 120 minutes. The reaction was terminated by adding three volumes of cold ACN with IS. Samples were centrifuged and analyzed by UPLC-MS/MS.

#### Plasma protein binding and blood-to-plasma partitioning

Plasma protein binding experiments were carried out in mixed-gender human plasma, male Sprague-Dawley rat plasma, male CD-1 plasma, male Beagle dog plasma, and male Cynomolgus monkey plasma (BioIVT) using a Pierce Rapid Equilibration Dialysis (RED) device. Briefly, plasma (300 μL), containing GS-441524 (10 μM), was loaded into two wells of the 96-well dialysis plate. Blank phosphate-buffered saline (PBS) (500 μL) was added to each corresponding receiver chamber. After 4 h of incubation (with shaking at 37°C), both sides were aliquoted, and matrix was normalized. Samples were centrifuged at 1000g for 10 mins after adding two volumes of ACN with IS. Supernatants were diluted 1:1 with water and analyzed with LC-MS/MS.

Plasma Protein binding was calculated as follows:

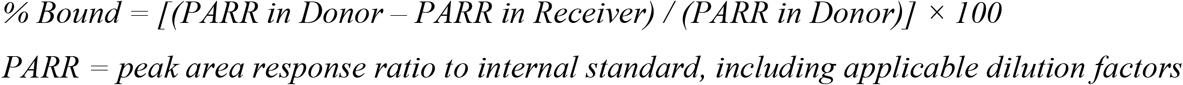

Blood partitioning studies were carried out in mixed-gender human whole blood, male Sprague-Dawley rat whole blood, male CD-1 whole blood, male Beagle dog whole blood, and male Cynomolgus monkey whole blood (BioIVT). Control plasma was obtained by centrifuging whole blood at 1000 g for 10 mins. Aliquots of whole blood and control plasma were then spiked with GS-441524 (10 μM) and incubated at 37°C for 1 h in a shaking water bath. The incubated whole blood was then centrifuged for 10 mins at 1000 g and plasma isolated from blood was separated. All samples of control plasma and plasma isolated from blood (blood plasma) were treated with four volumes of ACN with IS and centrifuged at 3000 rpm for 10 mins at 4°C. The supernatants were diluted 1:1 with water and analyzed by UPLC-MS/MS. The RBC to Plasma Partitioning (KRBC/P) and Whole Blood to Plasma Partitioning (KWB/P) were calculated as follows:

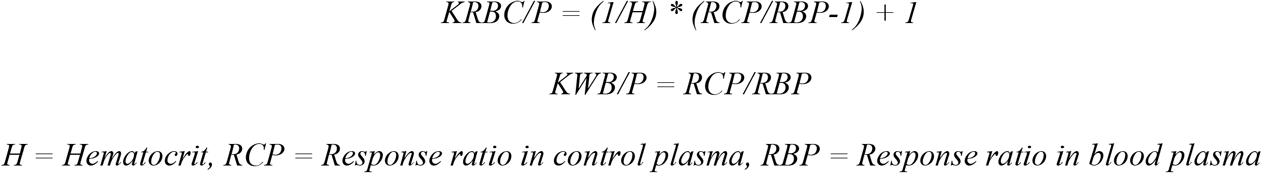

#### Metabolite identification in mouse, rat, dog, monkey, and human hepatocytes

Plateable cryopreserved hepatocytes from male CD-1 Mouse (Lot# DYF, 81.2% viability), male SD Rat (Lot# PMD, 88.1% viability), male Beagle Dog (Lot# YTK, 90.0% viability), male Cynomolgus Monkey (Lot# GJV, 88.7% viability), and male Human (Lot# PDC, 82.9% viability) were purchased from BioIVT (Baltimore, MD). Following vendors instructions, the hepatocyte vials were thawed in 37°C water bath and cells were suspended in 5 mL of corresponding plating media (antibiotic added, rodent media for mouse and rat, dog media for dog, and primate media for monkey and human). The cells were manually counted using Trypan Blue method and were diluted to target concentration (0.35 million/mL for mouse 0.7 million/mL for other species) with plating media. Diluted cells (500 µL) were then added to Collagen-I coated 24-well plate, distributed evenly, and incubated for 4 h for attaching. Media was changed media at 4 h and incubated overnight for compound incubations.

GS-441524 was incubated at 1 and 10 µM at 37°C for 24 h and 48 h with 5% CO2. After 24 h and 48 h, the reaction was stopped by adding 3x volumes of cold acetonitrile to the reaction mixture. Samples were vortexed and centrifuged to precipitate proteins. The supernatants were transferred to clean tubes and dried under a stream of N2 for ∼20 min at ambient temperature to remove most of organic solvent. The remaining liquid was dried by lyophilization. The residues were reconstituted with 10% methanol and injected onto the LC-UV/MS system for metabolite identification and profiling.

The analytical system consisted of a Dionex UHPLC 3000 (pumps, autosampler and PDA) interfaced to Orbitrap-Elite mass spectrometer. Liquid chromatography was accomplished using a Waters Atlantis T3 (5µm, 4.6×250 mm) column and a Phenomenex C18 (4×2mm) pre-column. Mobile phase consisted of 0.1% formic acid in water (solvent A) and 0.1% formic acid in water (solvent B) at 800 μL/min flow rate. The run started with 100% of solvent A for 5 minutes and gradually changed to 90% A after 25 minutes followed by 5% A from 28 to 30 minutes and 100% A from 30.5 to 40 minutes.

The Orbitrap-Elite mass spectrometer operated with a full scan range of m/z 100 – 1000 with resolution of 15000 and data dependent MS^n^ (n = 4) acquisition mode for analysis. The source operated in positive ionization mode with spray voltage of 4kV; capillary temperature, 320 °C; sheath gas flow rate 50 (arbitrary unit); auxiliary gas flow rate 15 (arbitrary unit); activation Q, 0.25; and collision energy, 35.

#### Transporter assays

##### MDR1 and BCRP substrate screening

Caco-2 (American Type Culture Collection), MDR1-MDCK (NIH), and BCRP-MDCK (transfected by Absorption Systems) cell monolayers were grown to confluence on collagen-coated, microporous membranes in 12-well assay plates. Caco-2 cells were cultured for 27 days, MDCK-MDR1 and MDCK-BCRP cells were cultured for 8 days prior to experiments. Two marker compounds, propranolol and atenolol, were used to confirm the integrity and function of the cell membrane. The experiment was carried out in Hanks’ balanced salt solution (HBSS) containing 10 mM HEPES and 15 mM glucose at pH 7.4. HBSS in the receiver chamber also contained 1% bovine serum albumin. Cells were first pre-incubated for 0.5 h in HBSS with or without the specific inhibitor (1 µM valspodar or 0.5 µM Ko143 for MDR1 and BCRP, respectively). The dosing solution contained 5 µM of GS-441524 with the presence or absence of the specific inhibitor. Cell monolayers were dosed on the apical side (A to B) or basolateral side (B to A) and incubated at 37°C with 5% CO_2_ in a humidified incubator. Samples were taken from the donor and receiver chambers at 2 h. Each experiment was performed in duplicate. All samples were assayed by LC-MS/MS. The apparent permeability (P_app_) and percent recovery were calculated as described previously [51].

##### Nucleoside transporters

MDCK-II cells were maintained in DMEM with low glucose and 10% FBS. Cells passages up to 40 were seeded at 60,000±10,000 cells/well on 96-well transwell membrane plates 24 h before transfection. Transporter assays were carried out 48 h after transfection. 96-well cell culture plate containing a monolayer of MDCK-II cells were grown on a permeable support and a corresponding 96-well receiver tray. Cell plates were maintained at 37°C in 5% CO2 atmosphere prior to initiation of the experiment. Experiments were conducted in HBSS for CNTs and ENT 1 and 2 or HBSS-Bis-Tris (pH 6.0) for ENT4. Each condition was run in triplicate. A volume of 150 µL buffer containing 10 µM, 30 µM (with or without reference inhibitor for each transporter), and 100 µM GS-441524 (0.5% DMSO) was added to basal compartment for ENT2, ENT4 assay. A volume of 100 µL buffer was added to apical compartment for ENT1 and CNTs assay. Plates were incubated at 37°C with orbital shaking at approximately 60 rpm for 5 min. The cells were washed on both the apical and the basolateral side for four times with ice cold PBS followed by addition of 60 µL cell extraction solution to the 96-well plate. Identical experiments were conducted using cells expressing the transporter of interest and control cells which did not express the transporter. The plates were agitated to mix the extract for 15 minutes on an orbital shaker at 120 rpm. A 30 µL sample extract was collected and mixed with 30 µL of internal standard before freezing the samples for further LC-MS/MS analysis. Activity was calculated by dividing the mean cellular accumulation in cells expressing the transporter by the mean cellular accumulation in control cells. The net transporter-mediated uptake rate of the substrate by each transporter was calculated:

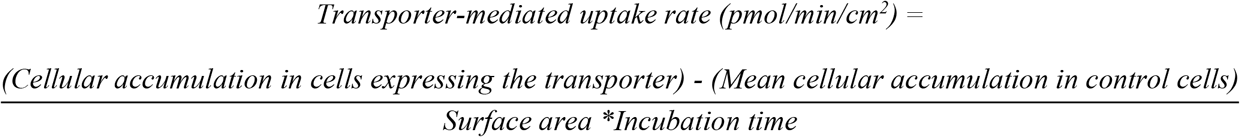

Percent inhibition was calculated by dividing the transporter-mediated uptake rate in presence of the test article or the reference inhibitor by the transporter-mediated uptake rate in presence of vehicle control:

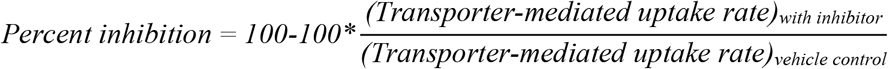

### Pharmacokinetics studies

Adult male C57BL/6 mice and Sprague Dawley rats were purchased from Charles River Laboratory (Wilmington, MA). PK studies for mice and rats were conducted at NIH animal facility (Bethesda, MD). In-life dog and monkey PK studies were conducted at Absorption System (San Diego, CA) and Frontage Labs (Concord, OH), respectively. Dogs and monkeys were fasted overnight prior to dose administrations. All experimental procedures were reviewed and approved by the Animal Care and Use Committee (ACUC) of the NIH Division of Veterinary Resources (DVR) or CRO labs for in-life studies.

Male C57BL6 mice (n = 3 per time point, 24 - 40 g) were dosed at 5 mg/kg IV or 10 mg/kg oral gavage. The dosing volume was 3 mL/kg for IV and 10 mL/kg for PO. All dosing solutions were freshly prepared on the day of administration. Blood samples were collected over 24 h.

Male Sprague–Dawley rats (n = 3 or 4 per treatment group, ∼ 343–375 g) were dosed in IV and PO dosing groups: 1 and 5 mg/kg IV, and 10, 30 and 100 mg/kg PO, respectively. Double cannulated rats (one catheter for IV dosing and another one for sampling) were used. Plasma samples were collected from jugular vein over 24 h. In bile duct cannulated (BDC) rat group, a single IV dose of 2 mg/kg was administered to three rats. Plasma, urine, and bile samples were collected over 48 h for BDC rats.

Male Cynomolgus monkeys (n = 3 per treatment group, 3.3–4.3 kg) received IV and PO administration of 2 and 5 mg/kg of GS-441524, respectively. The dosing volumes were 2 mL/kg for IV and 5 mL/kg for PO. Blood samples were obtained from each monkey up to 48 h after IV and PO administration. Urine samples were collected over 48 h for IV group.

Male Beagle dogs (n = 3 per treatment group, 9.0–11.4 kg) received 2 mg/kg IV and 5 mg/kg PO of GS-441524 administration. The dosing volume was 1.0 mL/kg for both groups. Plasma (both IV and PO groups) and urine (IV group) samples were collected over a period of 48 h.

All GS-441524 dosing solutions were prepared in 5% ethanol, 30% PG, 45% PEG-400 and 20% water + 1 equivalent of HCl. Blood samples from all groups and species were collected in K_2_EDTA tubes and stored on wet ice pending processing. Blood was centrifuged at 2200g at 4°C for 10 min to isolate plasma. All samples were stored at -80°C until analysis.

### Pharmacokinetics analysis

Phoenix WinNonlin, version 8.1 (Certara, St. Louis, MO), was used to perform PK analysis with the non-compartmental approach (Models 200 and 201 for PO and IV datasets). The area under the plasma concentration vs time curve (AUC) was calculated using the linear trapezoidal method. The terminal rate constant (λ) was obtained from the slope of at least three data points of the apparent terminal phase of the log-linear plasma concentration vs time curve when available. AUC0–∞ was estimated as the sum of the AUC0–t (where t is the time of the last measurable concentration) and Ct/λ. The apparent terminal half-life (t1/2) was calculated as 0.693/λ. Bioavailability (F) was calculated using ratios of dose normalized AUC (AUC/Dose) obtained from PO and IV PK studies.

Allometric methods were used to predict human plasma clearance and volume of distribution [21, 52]. Human PK prediction was carried out with GastroPlus, version 9.6 (SimulationsPlus, Lancaster, CA)

### UPLC-MS/MS bioanalysis

Ultra-performance liquid chromatography-tandem mass spectrometry (UPLC-MS/MS) methods were developed and optimized to determine GS-441524 concentrations in plasma, urine, and bile samples. The UPLC column was an Atlantis™ PREMIER BEH C18 AX (1.7 µm, 2.1×50 mm) purchased from Waters Corp. (Waltham, MA). The mobile phase A was 0.1% formic acid in water and B was 0.1% formic acid in acetonitrile. A gradient was run as follows: 0-0.2 min 1% B; 1.2 min 40% B; 1.3 min 95% B; 2.0 min 98% B; 2.1 min 1% B at a flow rate of 0.5 mL/min.

Mass spectrometric analysis was performed on a Waters Xevo TQ-S triple quadrupole instrument using electrospray ionization in positive mode with the selected reaction monitoring. The data was acquired and analyzed using TargetLynx XS version 4.1 (Waters Corp.). Positive electrospray ionization data were acquired using multiple reaction monitoring (MRM). The source settings for temperature, ion spray voltage and cone voltage were 600°C, 1.0 kV and 25 V, respectively. The MRM transitions were 292 →163, 292 →202 and 292 →147 for GS-441524 and 297→164 for SIL-IS.

The calibration standards and quality control samples (QCs) were prepared in the drug-free biological matrices. Pre-dose bile and urine samples were analyzed to confirm the absence of GS-441524 and used to prepare calibration standards and QCs. GS-441524 was extracted from plasma, bile, and urine samples by acetonitrile. Samples were prepared by adding a 20.0 μL aliquot of samples, QCs, and standards into the appropriate wells of a 96-well plate. A 200 μL aliquot of 50 ng/mL SIL-GS-441524 (internal standard, SIL-IS) in acetonitrile solution was added to the samples. All samples were then thoroughly shaken for 5 min. The plates were centrifuged at 3000 rpm and 4°C for 30 min and 150 μL supernatant was transferred to a 96-well plate. A 0.3 μL aliquot of plasma, urine, and bile supernatant were injected for UPLC-MS/MS analysis. The total run time was 3 min per sample.

For plasma and bile samples, the calibration standard for GS-441524 ranged from 1.00 to 10000 ng/mL. Quality control samples (QC, n=6) were prepared at 1.00 ng/mL, 10.0 ng/mL, 1000 ng/mL, and 10000 ng/mL in the corresponding matrix. The calibration standard ranged from 50.0 to 100000 ng/mL for urine samples. QCs (n=6) were prepared at 100 ng/mL, 1000 ng/mL, 10000 ng/mL, and 100000 ng/mL in the corresponding urine matrix. GS-441524 was quantified by calculating peak area ratios of GS-441524 to SIL-IS. Concentrations of GS-441524 were determined by a weighted (1/concentration^2^) least squares linear regression method.

## Acknowledgements

Authors would like to thank Dr. Zhiji Luo for his help in providing GS-441524 for all studies, Ms. Nida Delbe and Ms. Danitra Brathwaite of NIH DVR for their contributions to the in-life pharmacokinetic studies in rodents. We also like to thank Absorption Systems, Cyprotex PLC, Frontage laboratories and BioIVT for their contributions to selected in vitro ADME and PK studies, and Gilead Sciences for providing formulation protocol.

## Author contributions

A.Q.W. designed and performed experiments, analyzed data, interpreted results, and wrote the manuscript. N.R.H. performed experiments, analyzed data, and contributed to manuscript revision. E.C.P. performed experiments and simulation, interpreted results and contributed to writing. M.Y. performed simulation, interpreted results and contributed to writing. P.S. performed experiments and simulation, analyzed data, and contributed to writing. C.Z.C. designed experiments, interpreted results, and contributed to manuscript revision. W.H. provided the test compound and contributed to manuscript revision. P.T. contributed to manuscript revision. P.S. contributed to scientific discussion and manuscript revision. W.Z. designed experiments and contributed to writing. X.X. designed experiments, interpreted results, edited the manuscript, and acted as senior author of the study.

## REFERENCES

1. Warren, T.K., et al., Therapeutic efficacy of the small molecule GS-5734 against Ebola virus in rhesus monkeys. Nature, 2016. 531(7594): p. 381–5.

2. Beigel, J.H., et al., Remdesivir for the Treatment of Covid-19 - Final Report. N Engl J Med, 2020. 383(19): p. 1813–1826.

3. Gottlieb, R.L., et al., Early Remdesivir to Prevent Progression to Severe Covid-19 in Outpatients. N Engl J Med, 2021.

4. Beigel, J.H., et al., Remdesivir for the Treatment of Covid-19 — Final Report. New England Journal of Medicine, 2020. 383(19): p. 1813–1826.

5. Goldman, J.D., et al., Remdesivir for 5 or 10 Days in Patients with Severe Covid-19. N Engl J Med, 2020. 383(19): p. 1827–1837.

6. Owen, D.R., et al., An oral SARS-CoV-2 M(pro) inhibitor clinical candidate for the treatment of COVID-19. Science, 2021. 374(6575): p. 1586–1593.

7. Sheahan, T.P., et al., An orally bioavailable broad-spectrum antiviral inhibits SARS-CoV-2 in human airway epithelial cell cultures and multiple coronaviruses in mice. Sci Transl Med, 2020. 12(541).

8. Siegel, D., et al., Discovery and Synthesis of a Phosphoramidate Prodrug of a Pyrrolo[2,1-f][triazin-4-amino] Adenine C-Nucleoside (GS-5734) for the Treatment of Ebola and Emerging Viruses. Journal of medicinal chemistry, 2017. 60(5): p. 1648–1661.

9. Mulangu, S., et al., A Randomized, Controlled Trial of Ebola Virus Disease Therapeutics. New England Journal of Medicine, 2019. 381(24): p. 2293–2303.

10. Dickinson, P.J., et al., Antiviral treatment using the adenosine nucleoside analogue GS-441524 in cats with clinically diagnosed neurological feline infectious peritonitis. Journal of veterinary internal medicine, 2020. 34(4): p. 1587–1593.

11. Rasmussen, H.B., et al., Cellular Uptake and Intracellular Phosphorylation of GS-441524: Implications for Its Effectiveness against COVID-19. Viruses, 2021. 13(7).

12. Wei, D., et al., Potency and pharmacokinetics of GS-441524 derivatives against SARS-CoV-2. Bioorg Med Chem, 2021. 46: p. 116364.

13. Li, Y., et al., Remdesivir Metabolite GS-441524 Effectively Inhibits SARS-CoV-2 Infection in Mouse Models. J Med Chem, 2021.

14. Schafer, A., et al., Therapeutic efficacy of an oral nucleoside analog of remdesivir against SARS-CoV-2 pathogenesis in mice. bioRxiv, 2021.

15. Cox, R.M., et al., Oral prodrug of remdesivir parent GS-441524 is efficacious against SARS-CoV-2 in ferrets. Nat Commun, 2021. 12(1): p. 6415.

16. Warren, T.K., et al., Therapeutic efficacy of the small molecule GS-5734 against Ebola virus in rhesus monkeys. Nature, 2016. 531(7594): p. 381–385.

17. Tempestilli, M., et al., Pharmacokinetics of remdesivir and GS-441524 in two critically ill patients who recovered from COVID-19. Journal of Antimicrobial Chemotherapy, 2020. 75(10): p. 2977–2980.

18. Humeniuk, R., et al., Safety, Tolerability, and Pharmacokinetics of Remdesivir, An Antiviral for Treatment of COVID-19, in Healthy Subjects. Clinical and Translational Science, 2020. 13(5): p. 896–906.

19. Gorshkov, K., et al., The SARS-CoV-2 Cytopathic Effect Is Blocked by Lysosome Alkalizing Small Molecules. ACS Infect Dis, 2020.

20. In Vitro Drug Interaction Studies — Cytochrome P450 Enzyme- and Transporter-Mediated Drug Interactions, U.S.F.a.D. Administration, Editor. 2020.

21. Mahmood, I. and J.D. Balian, Interspecies scaling: predicting clearance of drugs in humans. Three different approaches. Xenobiotica, 1996. 26(9): p. 887–95.

22. Pruijssers, A.J., et al., Remdesivir Inhibits SARS-CoV-2 in Human Lung Cells and Chimeric SARS-CoV Expressing the SARS-CoV-2 RNA Polymerase in Mice. Cell Rep, 2020. 32(3): p. 107940.

23. Schooley, R.T., et al., Rethinking Remdesivir: Synthesis of Lipid Prodrugs that Substantially Enhance Anti-Coronavirus Activity. bioRxiv, 2020.

24. Yan, V.C. and F.L. Muller, Advantages of the Parent Nucleoside GS-441524 over Remdesivir for Covid-19 Treatment. ACS Med Chem Lett, 2020. 11(7): p. 1361–1366.

25. Ray, A.S., et al., Effective metabolism and long intracellular half life of the anti-hepatitis B agent adefovir in hepatic cells. Biochem Pharmacol, 2004. 68(9): p. 1825–31.

26. U.S. Food and Drug Administration website. Drugs@FDA:VEKLURY, Labeling-Package Insert (ORIG-1,10/22/2020). https://www.accessdata.fda.gov/drugsatfda_docs/label/2020/214787Orig1s000lbl.pdf.

27. Miller, S.R., et al., Remdesivir and EIDD-1931 Interact with Human Equilibrative Nucleoside Transporters 1 and 2: Implications for Reaching SARS-CoV-2 Viral Sanctuary Sites. Mol Pharmacol, 2021.

28. Human Protein Atlas 2021-02-24 [cited 2021 2021-11-4]; Available from: http://www.proteinatlas.org.

29. Administration, U.S.F.a.D., In Vitro Drug Interaction Studies — Cytochrome P450 Enzyme- and Transporter-Mediated Drug Interactions. 2020.

30. Katoh, M., et al., Kinetic analyses for species differences in P-glycoprotein-mediated drug transport. J Pharm Sci, 2006. 95(12): p. 2673–83.

31. Xia, C.Q., et al., Comparison of species differences of P-glycoproteins in beagle dog, rhesus monkey, and human using Atpase activity assays. Mol Pharm, 2006. 3(1): p. 78–86.

32. Li, M., et al., Identification of interspecies difference in efflux transporters of hepatocytes from dog, rat, monkey and human. Eur J Pharm Sci, 2008. 35(1-2): p. 114–26.

33. Mai, Y., et al., Quantification of P-Glycoprotein in the Gastrointestinal Tract of Humans and Rodents: Methodology, Gut Region, Sex, and Species Matter. Mol Pharm, 2021. 18(5): p. 1895–1904.

34. Uhlen, M., et al., Proteomics. Tissue-based map of the human proteome. Science, 2015. 347(6220): p. 1260419.

35. Lee, E.W., et al., Identification of the mitochondrial targeting signal of the human equilibrative nucleoside transporter 1 (hENT1): implications for interspecies differences in mitochondrial toxicity of fialuridine. J Biol Chem, 2006. 281(24): p. 16700–6.

36. Naes, S.M., et al., Equilibrative Nucleoside Transporter 2: Properties and Physiological Roles. Biomed Res Int, 2020. 2020: p. 5197626.

37. Owen, R.P., I. Badagnani, and K.M. Giacomini, Molecular determinants of specificity for synthetic nucleoside analogs in the concentrative nucleoside transporter, CNT2. J Biol Chem, 2006. 281(36): p. 26675–82.

38. King, J.R., et al., Drug-Drug Interactions between Sofosbuvir and Ombitasvir-Paritaprevir-Ritonavir with or without Dasabuvir. Antimicrob Agents Chemother, 2016. 60(2): p. 855–61.

39. Takahashi, K., et al., Contribution of equilibrative nucleoside transporter(s) to intestinal basolateral and apical transports of anticancer trifluridine. Biopharm Drug Dispos, 2018. 39(1): p. 38–46.

40. Martin, C., et al., High adenosine plasma concentration as a prognostic index for outcome in patients with septic shock. Critical Care Medicine, 2000. 28(9): p. 3198–3202.

41. Eltzschig, H.K., et al., HIF-1-dependent repression of equilibrative nucleoside transporter (ENT) in hypoxia. J Exp Med, 2005. 202(11): p. 1493–505.

42. Morote-Garcia, J.C., et al., Hypoxia-inducible factor-dependent repression of equilibrative nucleoside transporter 2 attenuates mucosal inflammation during intestinal hypoxia. Gastroenterology, 2009. 136(2): p. 607–18.

43. Lai, Y., C.M. Tse, and J.D. Unadkat, Mitochondrial expression of the human equilibrative nucleoside transporter 1 (hENT1) results in enhanced mitochondrial toxicity of antiviral drugs. J Biol Chem, 2004. 279(6): p. 4490–7.

44. Humeniuk, R., et al., Safety, Tolerability, and Pharmacokinetics of Remdesivir, An Antiviral for Treatment of COVID-19, in Healthy Subjects. Clin Transl Sci, 2020. 13(5): p. 896–906.

45. Chen, C.Z., et al., Drug Repurposing Screen for Compounds Inhibiting the Cytopathic Effect of SARS-CoV-2. Frontiers in Pharmacology, 2021. 11: p. 592737.

46. Reed, L.J. and H. Muench, A SIMPLE METHOD OF ESTIMATING FIFTY PER CENT ENDPOINTS12. American Journal of Epidemiology, 1938. 27(3): p. 493–497.

47. Sun, H., et al., Predictive models of aqueous solubility of organic compounds built on A large dataset of high integrity. Bioorganic & medicinal chemistry, 2019. 27(14): p. 3110–3114.

48. Sun, H., et al., Highly predictive and interpretable models for PAMPA permeability. Bioorganic & medicinal chemistry, 2017. 25(3): p. 1266–1276.

49. Shah, P., et al., Predicting liver cytosol stability of small molecules. Journal of cheminformatics, 2020. 12(1): p. 21.

50. Shah, P., et al., An Automated High-Throughput Metabolic Stability Assay Using an Integrated High-Resolution Accurate Mass Method and Automated Data Analysis Software. Drug metabolism and disposition: the biological fate of chemicals, 2016. 44(10): p. 1653–1661.

51. Hubatsch, I., E.G. Ragnarsson, and P. Artursson, Determination of drug permeability and prediction of drug absorption in Caco-2 monolayers. Nat Protoc, 2007. 2(9): p. 2111–9.

52. Sharma, V. and J.H. McNeill, To scale or not to scale: the principles of dose extrapolation. Br J Pharmacol, 2009. 157(6): p. 907–21.

